# Automated analysis of PD1 and PDL1 in lymph nodes and the microenvironment of transmissible tumors in Tasmanian devils

**DOI:** 10.1101/2022.10.31.513798

**Authors:** Grace G Russell, Chiara Palmieri, Jocelyn Darby, Gary P. Morris, Nicholas M. Fountain-Jones, Ruth J. Pye, Andrew S. Flies

## Abstract

The wild Tasmanian devil (*Sarchophilus harrisii*) population has suffered a devastating decline due to two clonal transmissible cancers. Devil facial tumor 1 (DFT1) was first observed in 1996, followed by a second genetically distinct transmissible tumor, devil facial tumor 2 (DFT2), in 2014. DFT1/2 frequently metastasize, with lymph nodes being common metastatic sites. Downregulation of MHC-I by DFT1 cells is a primary means of evading allograft immunity aimed at polymorphic MHC-I proteins. DFT2 cells constitutively express MHC-I, and MHC-I is upregulated on DFT1/2 cells by interferon gamma, suggesting other immune evasion mechanisms may contribute to overcoming allograft and anti-tumor immunity. Human clinical trials have demonstrated PD1/PDL1 blockade effectively treats patients showing increased expression of PD1 in tumor draining lymph nodes, and PDL1 on peritumoral immune cells and tumor cells. The effects of DFT1/2 on systemic immunity remain largely uncharacterized. This study applied the open-access software QuPath to develop a semiautomated pipeline for whole slide analysis of stained tissue sections to quantify PD1/PDL1 expression in devil lymph nodes. The QuPath protocol provided strong correlations to manual counting. PD-1 expression was approximately 10-fold higher than PD-L1 expression in lymph nodes and was primarily expressed in germinal centers, whereas PD-L1 expression was more widely distributed throughout the lymph nodes. The density of PD1 positive cells was increased in lymph nodes containing DFT2 metastases, compared to DFT1. This suggests PD1/PDL1 exploitation may contribute to the poorly immunogenic nature of transmissible tumors in some devils and could be targeted in therapeutic or prophylactic treatments.

## Introduction

Cancer is prevalent in nearly all mammalian species (1), but cancer immunology research has primarily focused on humans, mice, and a few other species (2). Naturally transmissible monoclonal cancers provide an opportunity to study cancer with the tumor genotype held relatively constant, whilst the host genotype and environment vary in a real-world cancer model. There are three known naturally transmissible cancers in mammals. The canine transmissible venereal tumor (CTVT) is a sexually transmitted tumor that has been circulating in dogs (*Canis lupus familiaris)* for an estimated 4,500 to 8,500 years (3,4). More recently, two different transmissible Tasmanian devil facial tumors (DFT1, DFT2) have emerged in wild Tasmanian devils (*Sarcophilus harrisii*) (**Figure 1)** (5,6). Both CTVT and DFT1 tumor cells have passed through thousands of individuals, and thus have successfully evaded the host immune system despite genetic differences between tumor and host cells.

**Figure 1.**
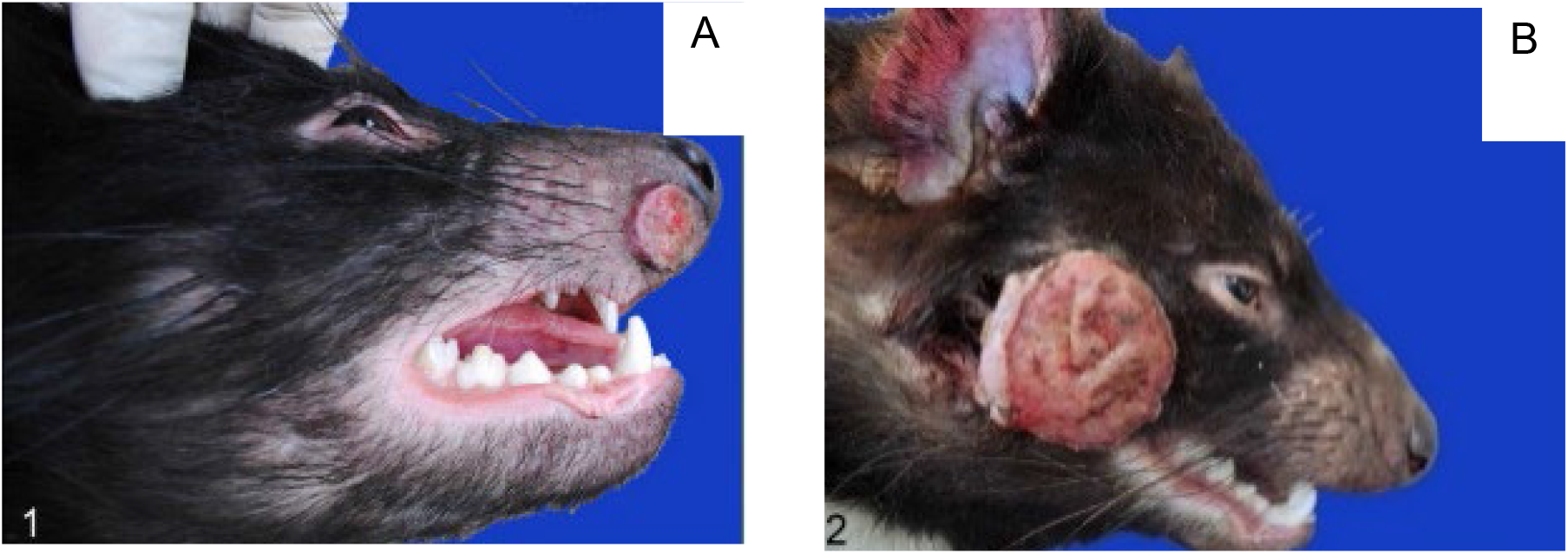
Devil facial tumors. Ulcerated (A) DFT1 and (B) DFT2 mass on devil faces.

DFT1 and DFT2 arise from Schwann cell-like progenitor cells (7,8). Both tumors commonly metastasize to the lymph nodes, lungs, liver, and kidneys and are nearly always fatal within 6- 12 months of first detection (6,9). DFT1 was first documented in 1996 and has been the driving factor behind the 80% decline in devil populations in the past 25 years. This has resulted in devils being placed on the endangered species list (10). Immunology research supports conservation efforts aimed at controlling devil facial tumor disease (DFTD; disease caused by either DFT1 or DFT2 cells) in devils, and provides an unprecedented opportunity to study a real-world, large animal cancer model.

A pivotal finding in determining the pathogenesis of DFT1 was the demonstration of genetic and epigenetic downregulation of MHC-I surface expression on DFT-1 cells (11). DFT1 cells harbor a hemizygous deletion in the β2 microglobulin (B2M) (12), which is part of the MHC-I antigen presentation complex and is critical to T cell immunity. This is hypothesized to play a key role in the absence of MHC-I on the surface of DFT1 cells. However, MHC-I can be upregulated following stimulation with interferon gamma (IFNG) (11,13,14). Subsequent studies identified MHC-I as a primary antibody target in immunotherapy-induced and rare cases of naturally occurring DFT1 tumor regression (15–18). By contrast, DFT2, which only emerged in 2014, constitutively expresses surface MHC-I (19), suggesting other immune evasion mechanisms are likely needed to overcome allograft and damage-induced anti-cancer immunity.

In the past decade, immune checkpoints have been shown to play a key role in regulating anti- cancer immunity in humans and mice. Multiple therapeutic antibodies targeting the PD1/PDL1 pathway have shown success in the treatment of human cancers (20,21). These antibodies neutralize the inhibitory effects of PDL1 and can rescue effector cells from PD1-induced immunosuppression and enhance anti-tumor immunity (22). However, the PD1 blockade has also proven effective in a minority of patients with low or no PDL1 expression on tumor cells (23). This has led to the discovery that both PD1 expression in tumor draining lymph nodes and PDL1 expression on tumor infiltrating immune cells, can provide prognostic information on the response to PD1/PDL1 inhibition, independent of PDL1 expression on tumor cells (24,25). Given these findings, a thorough investigation into the PD1/PDL1 pathway in devil lymphoid structures and the DFT1/2 microenvironment is needed.

We previously developed devil specific anti-PD1 and anti-PDL1 monoclonal antibodies and qualitatively showed that PD1 and PDL1 were expressed in devil lymph nodes (26). Furthermore, we have found that PDL1 can be upregulated on DFT1/2 cells in response to IFNG treatment (13,26). Interestingly, genetically modifying DFT1 cells to constitutively express NLRC5, a master regulator of MHC-I expression, results in constitutive surface expression of MHC-I but not PDL1 (18). However, there are no studies quantifying PD1 and PDL1 expression in lymph nodes or the metastatic tumor microenvironment of any marsupial species.

Automated image analysis software can generate both quantitative and spatial information about PDL1 and other immune checkpoints in the tumor microenvironment (27–29). For example, quantifying the degree of invasion of immune cells into tumors and identifying interactions between different cell types can provide useful information for understanding tumor microenvironments (30,31). QuPath (<u>Qu</u>antitative <u>Path</u>ology), an open-access whole slide analysis software, has been successfully used for automated quantitative image analysis in various fields of primary research and diagnostic pathology (32,33). Importantly, the use of QuPath has been shown to produce high quality, quantitative, reproducible, objective data when a validated image analysis pipeline is established for the project (34–36). Additionally, once automated analysis pipelines have been established, they can open up immunohistochemical (IHC) analytical techniques to researchers without specializations in pathology and provide digital records of IHC classifications.

In the present study we developed a QuPath pipeline to quantify PD1 and PDL1 expression in the DFT1/2 tumor microenvironment and in devil lymph nodes with and without tumor metastases. We validated this pipeline using Matthew’s correlation coefficient (MCC) and by calculating sensitivity, specificity, accuracy, and precision, showing that our automated pipeline produced highly specific and accurate quantification. Using this pipeline, we found that PD1 expression was high in lymph node germinal centers. We also observed restricted PDL1 expression in lymph nodes and the tumor microenvironment.

## Materials and Methods

### Tasmanian devil tissues and sample processing

DFT1 and DFT2 tumor sections and devil lymphoid tissue samples were sourced from archived formalin-fixed paraffin embedded (FFPE) samples at the Menzies Institute for Medical Research, University of Tasmania. Samples were collected during necropsies from euthanized devils as listed in **Table S1**. One devil lymph node was sourced from a wild devil with no apparent tumors that was killed by a vehicle. The carcass was stored on ice for approximately 12 hours prior to necropsy. The three remaining healthy lymph node samples were provided by Natural Resources and Environment Tasmania sourced from captive devils. Lymph node samples were collected and fixed in 10% neutral-buffered formalin, before being placed in 70% ethanol and stored at room temperature until processing. Tasmanian devil tissue was collected under University of Tasmania Animal Ethics Committee permit numbers A0012513, A0014976, A0017469, A0017550. Paraffin embedding of tissue sections was performed using techniques similar to previous studies (26,37). Full details are available in **Supplementary Methods 1**.

### Immunohistochemical staining

FFPE tissue sections were deparaffinized and antigen retrieval was undertaken in 0.1 M citric acid buffer solution (Dako, target retrieval solution, S169984-2) using an electric pressure cooker (Russell Hobbs RHNHP401). This was followed by blocking the endogenous peroxidase activity with 10% hydrogen peroxide (H_2_O_2_) solution for 10 minutes. To prevent non-specific antibody binding, a commercial protein blocking solution (Dako, Serum Free, X0909) was applied to the slides and incubated for 30 min.

Tissue sections were then incubated with anti-PD1 (1:300, 3G8) and anti-PDL1 (1:100, 1F8) in antibody diluent (Dako, S0809) for 60 min (26). Slides were washed in PBS and incubated with horseradish peroxidase (HRP) conjugated secondary antibody included in the peroxidase substrate kit (Dako EnVision + System HRP labelled polymer anti-mouse or anti-rabbit IgG, K4001 and K4003), for 30 min at room temperature. Diaminobenzidine (DAB; Dako Liquid DAB+ Substrate Chromogen System K3436) was applied to each slide for 10 min and incubated at RT. Tissue sections were counterstained with Mayer’s hematoxylin (Thermo Fisher Scientific, FNN11008) followed by dehydration and mounting. Negative controls were achieved using mouse IgG1-κ isotype control (PharMingen, 557273) and a secondary only control.

### QuPath image analysis

Following immunohistochemical staining, slides were scanned using a VS120 Virtual Slide System (Olympus) at 40x magnification. Digitalized slides were saved in .vsi format. Digitalized slide images were imported into the open-source software QuPath, with the image type set to “Brightfield (H-DAB)” and the stain vectors automatically estimated. Each image was duplicated, to produce an image for nuclei detection/cell classification optimization, and final analysis. The analysis pipeline was modified from the QuPath online brightfield analysis tutorial v0.2.3 (38). Custom scripts for QuPath were developed with the aid of IntelliJ ® Community Edition v2020.3.2 using Apache Groovy coding language. All scripts are available in **Supplementary dataset 2**.

### Image analysis – nuclei detection optimization and supervised machine learning cell classification training

The QuPath *Cell detection* plugin, optimized for each image, was used to detect cell nuclei. Briefly, lymph node sections were annotated with 20 100×100 μm rectangle regions of interest (ROI), distributed throughout the tissue section (**Figure 2D**). Nuclei were manually counted using the *Points* function to provide a gold standard for the number and classification (DAB positive or DAB negative) of cells in each ROI (**Figure 2E**). Twenty-four *Cell detection* scripts with varying parameter combinations were written and applied to all images in the project using the *batch* function in the QuPath *Script editor*. The numbers of cells detected in each ROI, in each image, for each *Cell detection* script, was then compared to the manual counts through a Pearson’s correlation. Strong cytoplasmic DAB staining impacted the ability of some scripts to identify the hematoxylin-stained nuclei. Each script was therefore assessed through both correlation and manual visual performance. Scripts that correlated significantly to manual counts, and which were visually performing well were selected for further use.

**Figure 2.**
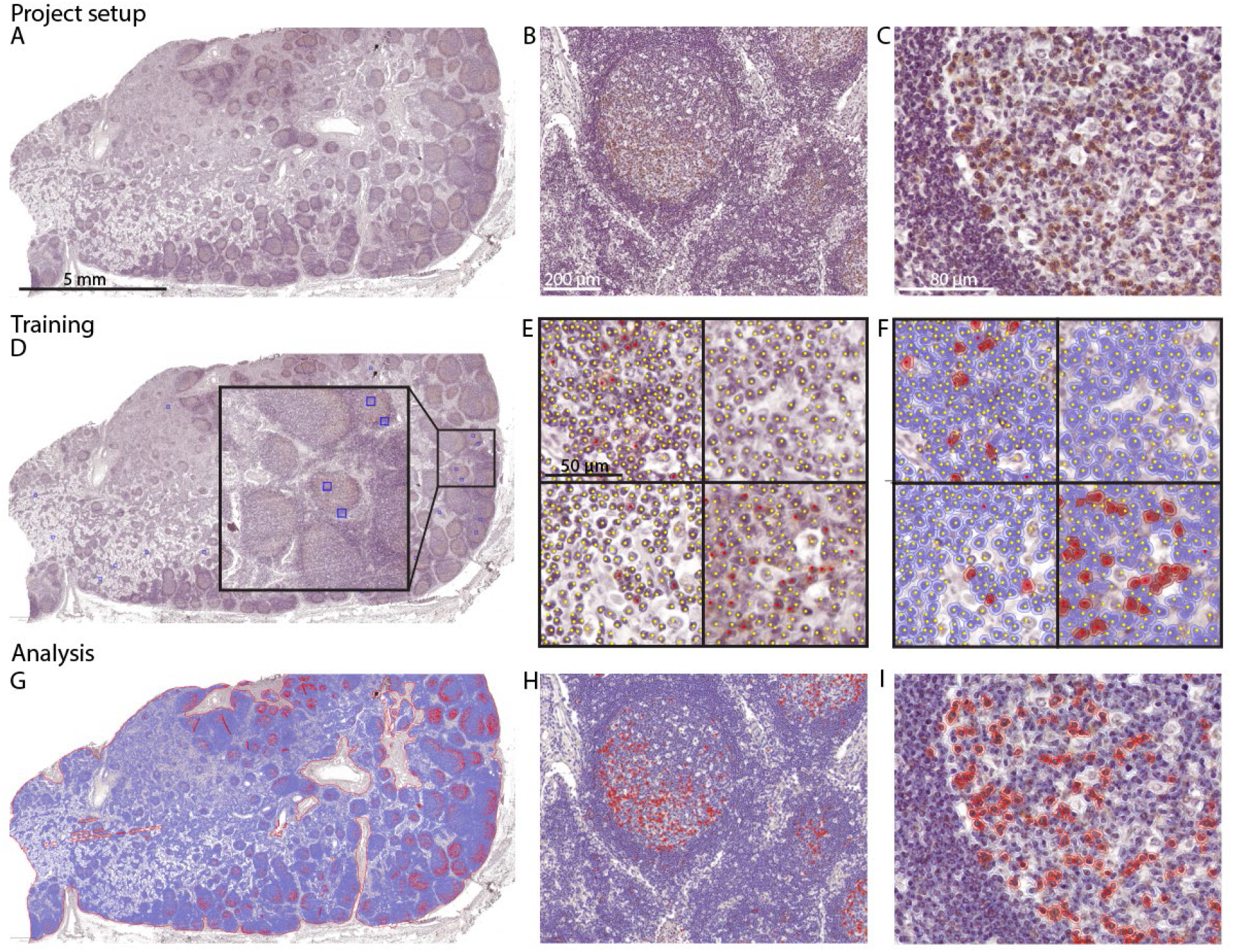
QuPath pipeline and image analysis. (A-C) Tissue samples are scanned using whole slide imaging and data files are uploaded into QuPath. (D) 100×100 µm regions of interest (ROI; blue squares) representative of a range of tissue staining characteristics and anatomical locations were selected throughout the lymph node. (E) Cells were annotated as positive (red) or negative (yellow) and (F) these annotations used to optimize cell detection, and train machine learning cell classifiers. (G-I) Tissue sections to be analyzed were then annotated and the optimized cell detection and classification systems applied (G-I), followed by data export. Images shown are from TD504 submandibular lymph node stained with anti-PD1 (3G8).

The QuPath trainable *Object classifier* plugin, using an inbuilt random forest machine learning algorithm, was used to classify cells as DAB positive or DAB negative. An individualized classifier for each image was created to eliminate the effects of inter- and intra-batch staining variation (31). Cells that had been annotated as positive or negative during nuclei detection optimization were used as training datasets for each *Object classifier*. The optimized *Cell detection* script for each image was run, followed by object classifier training. To avoid any substantial impact of non-specific nuclear background staining on the final analysis, ROI used for training machine learning classifiers were intentionally placed over areas where background noise was present, and the classifiers were then trained to recognize the background noise as negative.

The performance of each classifier was assessed through sensitivity, specificity, precision, accuracy, and the MCC. The 20 ROI were randomly divided into 16 training ROI and 4 testing ROI. Training was then performed on the 16 training ROI and the outcomes of the validation ROI were used to produce confusion matrices and MCC for each image.

### Automated Image Analysis

Analysis of anti-PD1 and anti-PDL1 tissue cohorts was performed using the optimized nuclear detection parameters and *Object classifier* for each image. Gross tissue detection was performed using the QuPath *Magic* wand function, and the *Brush* function to smooth edges and exclude tissue artefacts. The lymph node was manually annotated around its margin, inclusive of tumor metastases if they were present (**Figure 2G)**. Tumors were separately annotated using the *brush* function. Tumor metastases’ margins were identified as the region internal to a fibrous capsule of DFT1 metastases, or the margin between tumor cells and immune cells in DFT2 metastases. Following tissue annotation, the optimized nuclei detection algorithms and *Object classifiers* were applied to each image to detect nuclei and classify cells as DAB positive or negative (**Figure 2G & H)**. To provide a measure of positively labelled cell density (cells/mm^2^), data on the classification and number of cells in each tissue region (lymph node, tumor metastasis) along with the area (mm^2^) were exported and processed in Microsoft Excel and GraphPad Prism v9.0.

### Statistical analysis

Correlation measurements between manual and automated cell detection scripts were performed in GraphPad Prism v9.0. Correlation data assessed using the Pearson correlation coefficient.

Machine learning classifiers were analyzed using a confusion matrix (also known as a contingency matrix) (**Figure S3**) and MCC. A confusion matrix was produced using the true positive (TP), true negative (TN), false positive (FP), and false negative (FN) cell counts for each individual fold. The sensitivity, specificity, accuracy, precision, and negative predictive value along with an MCC were calculated for each individual confusion matrix.

MCC is a commonly used performance metric to assess the overall performance of machine learning classifiers when handling unbalanced binary datasets (39). MCC is calculated using the formula:

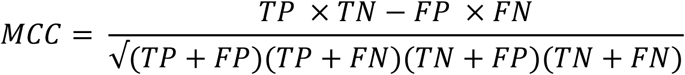

The data input from confusion matrices were used to calculate the MCC for binary comparisons. Results are produced in the range [-1, +1] where +1 is achieved in the case of perfect correlation, whereas -1 is achieved with perfect misclassification (e.g., all positives are classed as negative, and all negatives classed as positive). An MCC score of 0 can be expected to be produced by a classifier that randomly classifies points as positive or negative (40). The size of all correlations was interpreted as: 0 to 0.1 – negligible correlation; 0.1 to 0.39 – weak correlation; 0.4 to 0.69 – moderate correlation; 0.7 to 0.89 – strong correlation and 0.9 to 1.0 – very strong correlation (41).

All statistical analysis was performed in GraphPad Prism v9.0. Outlier testing was not conducted due to the limited number of biological samples. Due to the small number of datapoints, data were assessed for normality using QQ plots (42). Comparisons were between the density of positively labelled cells (cells/mm^2^) in DFT1, DFT2, and normal whole lymph node sections. Non-normally distributed data were log transformed prior to applying a one-way ANOVA. The density of PD1 or PDL1 labeled cells in lymph nodes containing DFT1 or DFT2 tumor metastases, and the density of PD1 or PDL1 labelled cells in DFT1 or DFT2 tumor metastases was compared using a two-way ANOVA without repeated measures, followed by Tukey’s multiple comparisons test. A p-value <0.05 was considered statistically significant.

## Results

### Variability of fully automated nuclei detection outcomes

We first aimed to develop a single nuclei detection classifier that could be used for every image in the project, as has been achieved in similar studies (31,33,43). Twenty-four nuclei detection scripts were trialed with differing optical density parameters (**supplementary dataset 2**). The script denoted V13 was found to be the most highly correlated algorithm to the manual counts across both anti-PD1 (3G8) and anti-PDL1 (1F8) cohorts with a Pearson’s correlation of r = 0.89 and r = 0.86, respectively (**Figure 3**). Further scrutiny of these data determined the range of correlations achieved for each image in the cohort to be 0.73-0.98 for the anti-PD1 (3G8) cohort, and 0.80-0.98 for the anti-PDL1 (1F8) cohort (**Figure 3**), suggesting high variability.

**Figure 3.**
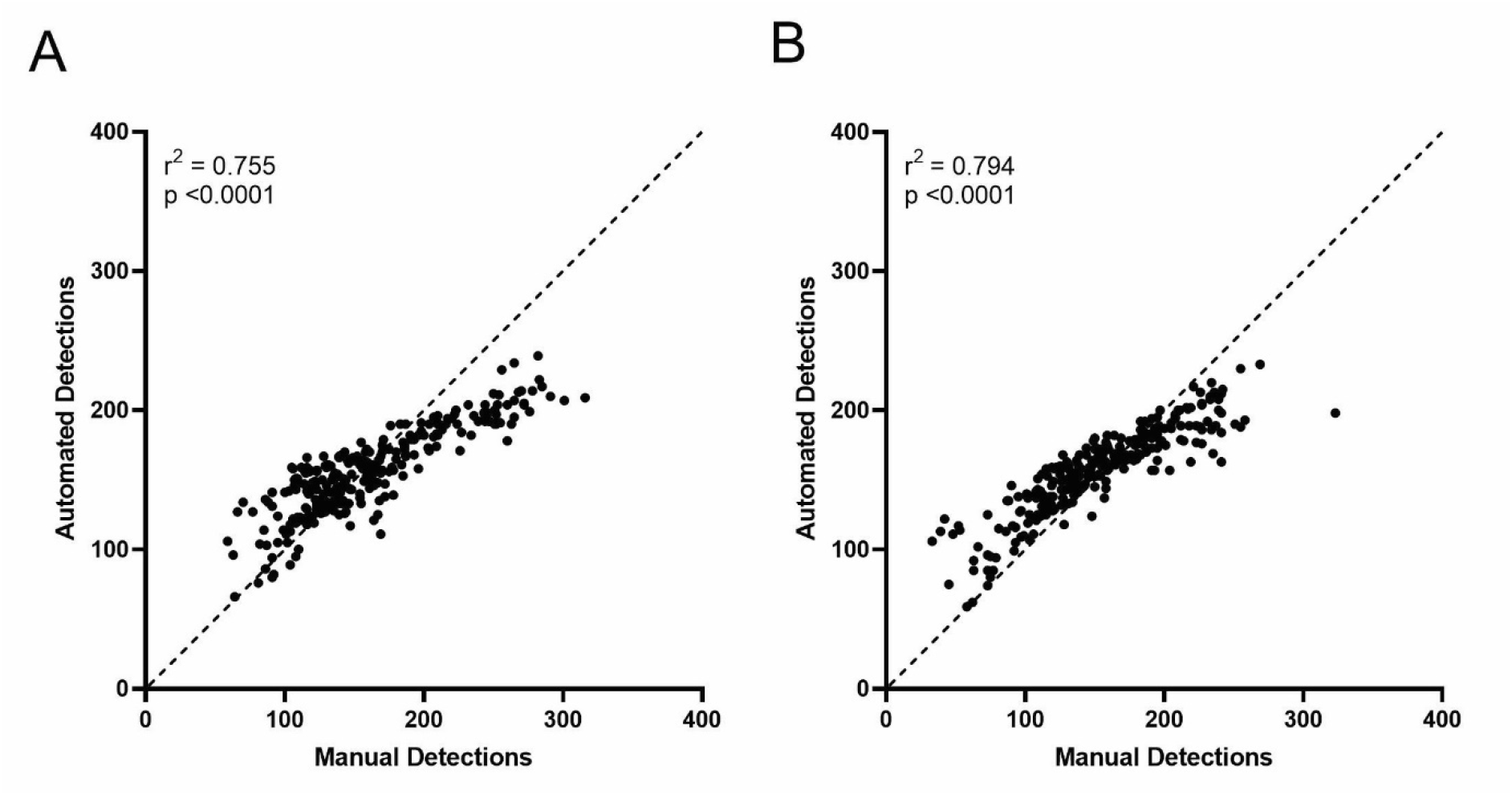
Correlations of manual and automated nuclei detections. Nuclei detection script V13 achieved the highest overall correlation when applied to the whole (A) anti- PD1 and (B) anti-PDL1 tissue cohorts, however the correlations achieved using this script on individual images showed marked variation, with some unacceptably low. Spearman correlation (r_s_) and *P-*values are shown.

These results demonstrated that it was not feasible to develop a single nuclei detection algorithm capable of correctly identifying nuclei for all images in our project. Additionally, some highly correlated scripts did not identify nuclei in cells with strong cytoplasmic DAB staining (i.e., false-negatives, **Figure S2)**. Characteristic features of DFTD tumors (vesicular nuclei with altered chromatin pattern) were also not consistently recognized by the nuclei detection algorithm which has been optimized to recognize lymphocyte nuclei. To overcome this issue, an additional visual assessment of the nuclei detection achieved with each script was performed. The final selection of scripts for each individual image was ultimately based off both the correlation data and a visual confirmation that the script was able to identify cells with strong DAB staining. The individual nuclei detection correlation coefficients for every image in the project are listed in **Table 1**.

**Table 1.**
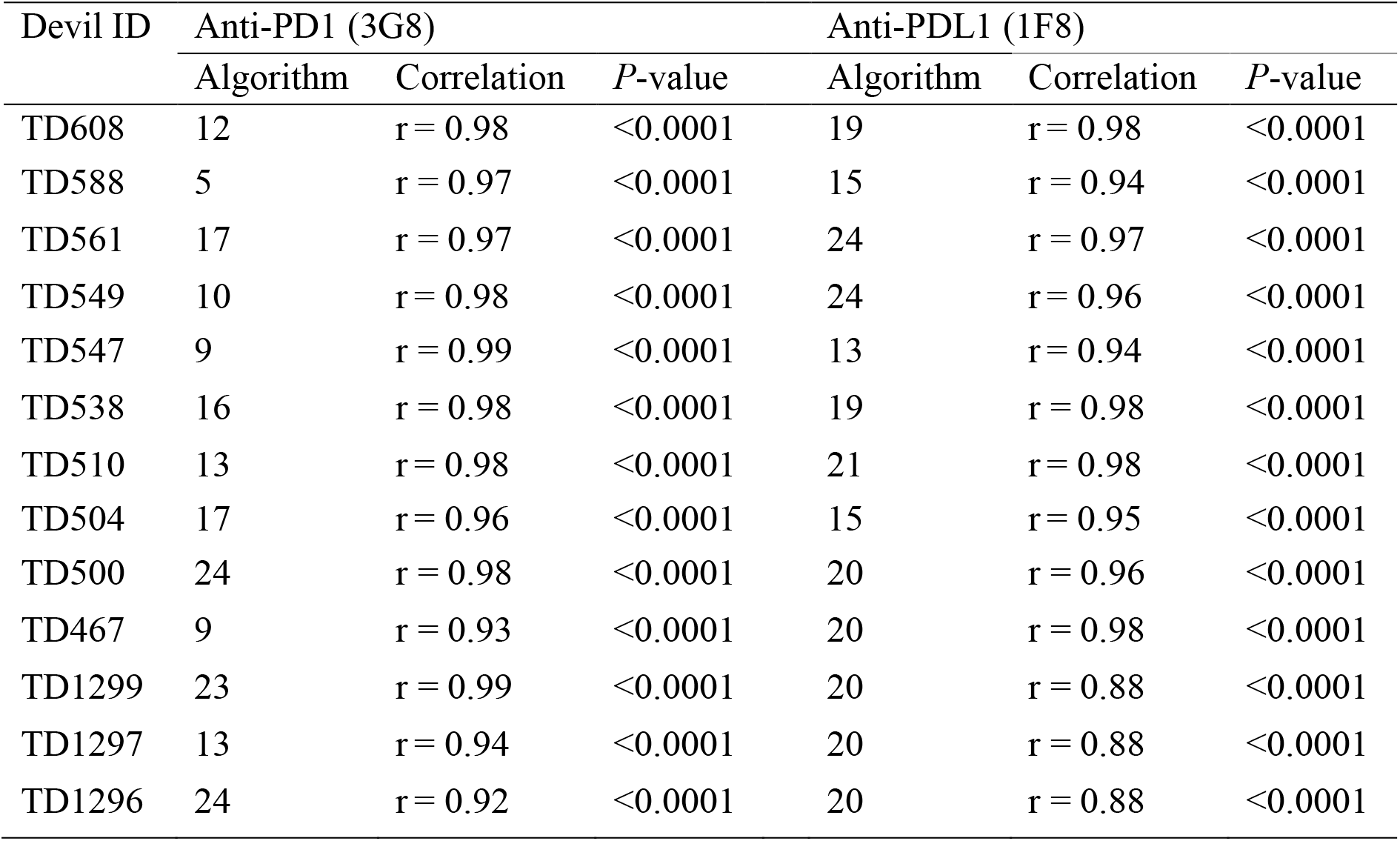
Pearson’s correlation between manual and optimized nuclei detection algorithms.

### Supervised machine learning classifiers produced accurate PD1 and PDL1 detection

Next, we assessed the performance of the supervised machine learning classifiers. These classifiers were trained to classify previously detected cells as DAB positive or negative based on the DAB staining within the cytoplasm. The classifiers exhibited a high specificity, precision, and accuracy. The MCC demonstrated the predictive performance of most classifiers was strong (MCC >0.7) to very strong (MCC>0.9) (**Table 2**). However, many of the trained classifiers underestimated the number of positively stained cells (high false negative), resulting in a sensitivity <70% in thirteen instances.

**Table 2.** Confusion matrix and MCC values of cell classification training. Outcomes of a randomly selected fold of training input used to develop the RTree algorithms used for classification of cells in anti-PD1 and anti-PDL1 cohorts.

| Devil ID | Sensitivity (%) | Specificity (%) | Precision (%) | Accuracy (%) | MCC |
| --- | --- | --- | --- | --- | --- |
| Anti-PD1 (3G8) |  |  |  |  |  |
| TD608 | 70 | 98 | 80 | 96 | 0.73 |
| TD588 | 86 | 99 | 93 | 98 | 0.89 |
| TD561 | 81 | 98 | 84 | 95 | 0.80 |
| TD549 | 98 | 99 | 87 | 99 | 0.92 |
| TD547 | 68 | 99 | 88 | 97 | 0.76 |
| TD538 | 44 | 100 | 89 | 98 | 0.62 |
| TD510 | 62 | 99 | 73 | 99 | 0.66 |
| TD504 | 70 | 99 | 82 | 96 | 0.74 |
| TD500 | 60 | 100 | 94 | 99 | 0.74 |
| TD467 | 68 | 100 | 93 | 98 | 0.79 |
| TD1299 | 63 | 100 | 92 | 99 | 0.76 |
| TD1297 | 75 | 100 | 100 | 100 | 0.87 |
| TD1296 | 54 | 99 | 87 | 95 | 0.66 |
| Anti-PLD1 (1F8) |  |  |  |  |  |
| TD608 | 100 | 100 | 100 | 100 | 1.00 |
| TD588 | 100 | 100 | 100 | 100 | 1.00 |
| TD561 | 67 | 100 | 67 | 100 | 0.66 |
| TD549 | 100 | 100 | 100 | 100 | 1.00 |
| TD547 | 80 | 100 | 100 | 100 | 0.89 |
| TD538 | 100 | 100 | 100 | 100 | 1.00 |
| TD510 | 100 | 100 | 100 | 100 | 1.00 |
| TD504 | 50 | 100 | 100 | 100 | 0.71 |
| TD500 | 50 | 100 | 100 | 100 | 0.71 |
| TD467 | 57 | 100 | 100 | 99 | 0.75 |
| TD1299 | 33 | 100 | 100 | 99 | 0.57 |
| TD1297 | 100 | 100 | 100 | 100 | 1.00 |
| TD1296 | 50 | 100 | 100 | 99 | 0.70 |

### Lymph node anatomy and staining

Thirteen lymph node sections were analyzed (4 healthy, 5 DFT1, 4 DFT2) and the density of positively labelled cells (cells/mm^2^) was compared between the three groups. Lymph node features including lymphatic follicles containing a germinal center and mantle zone, paracortex, and medulla were observed in all lymph nodes obtained from healthy, DFT1-, and DFT2-infected animals. The lymph nodes from healthy captive devils TD1296, TD1297 and TD1299 contained 9, 2, and 6 secondary follicles, respectively (**Figure 4, S3**). In comparison to the lymph node from TD608, the wild devil killed on the road but with no signs of DFT1 or DFT2, contained 39 secondary follicles. The average number of secondary follicles present in devils infected with DFT1 and DFT2 were 90 (range 20-147) and 78 (range 23-216), respectively (**Table S2**). A Kruskal-Wallis test demonstrated no significant difference (Kruskal-Wallis statistic = 4.711, p = 0.0324) between the number of secondary lymphatic follicles between the healthy, DFT1, and DFT2 cohorts (**Figure S3**).

**Figure 4.**
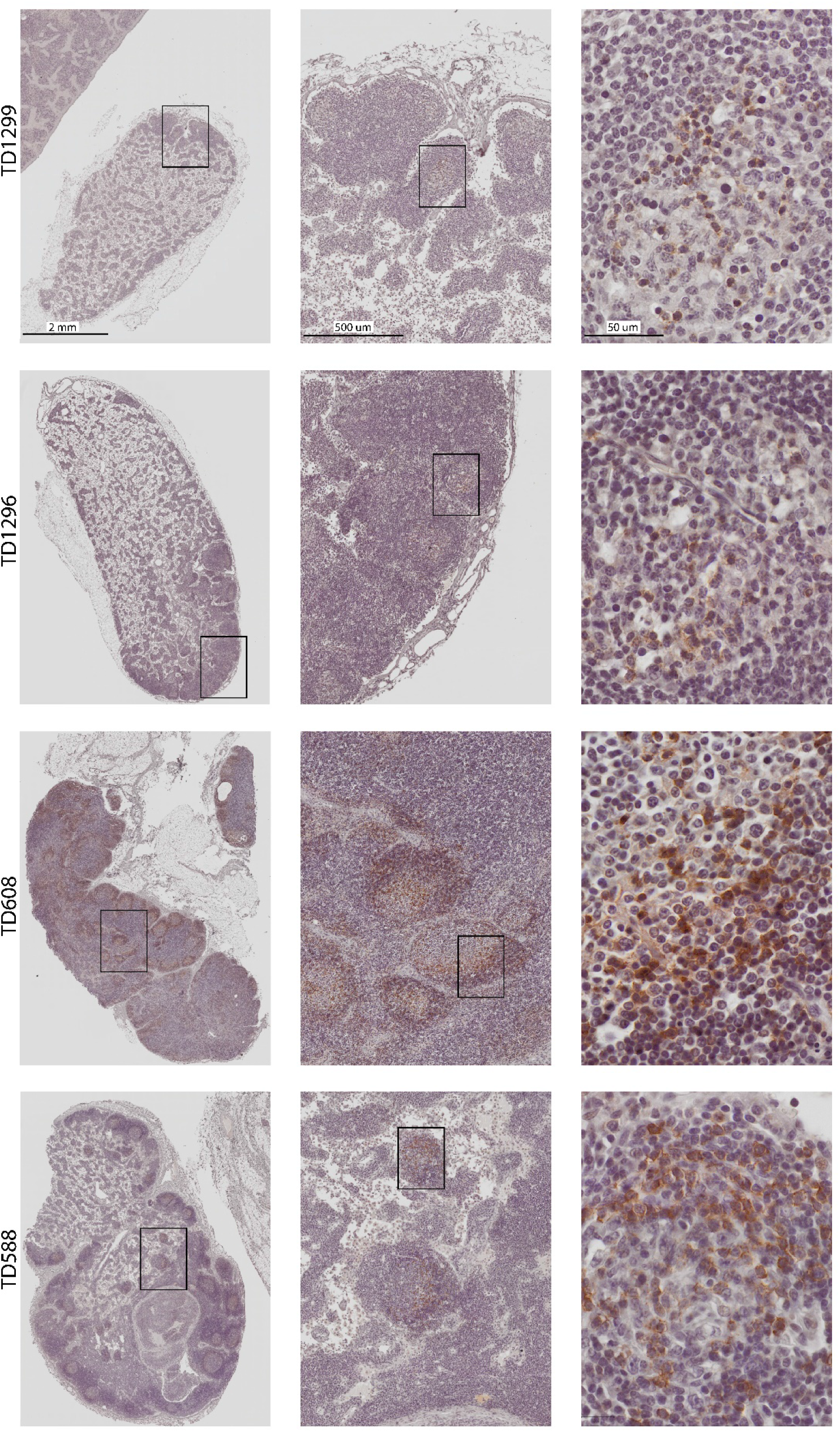
IHC showing PD1 labeled devil lymph nodes and tumor metastases. The submandibular lymph nodes from healthy captive devils (TD1296; TD1298), wild roadkill devil (TD608), and wild DFT1 infected devil (TD588) are shown. There was an increase in the number of secondary lymphoid follicles and density of positively labeled PD1 in all wild devils when compared to the healthy captive controls.

Moderate to strong PD1 labeling of lymphocyte cell membranes was identified in all lymph nodes (**Figure 4; Figure 5**). PD1 labeled cells were predominantly located within the germinal centers of secondary follicles and occasionally within the paracortex. PD1 labeled tumor infiltrating immune cells were infrequently identified in DFT1 and DFT2 tumor metastases, predominantly in the inner tumor margin and fibrous tumor stroma (**Figure 5)**. PD1 labeling in the lymph node of the wild healthy devil, TD608, was stronger, and PD1 labeled cells were more numerous than the other healthy control devils (**Figure 4**).

**Figure 5.**
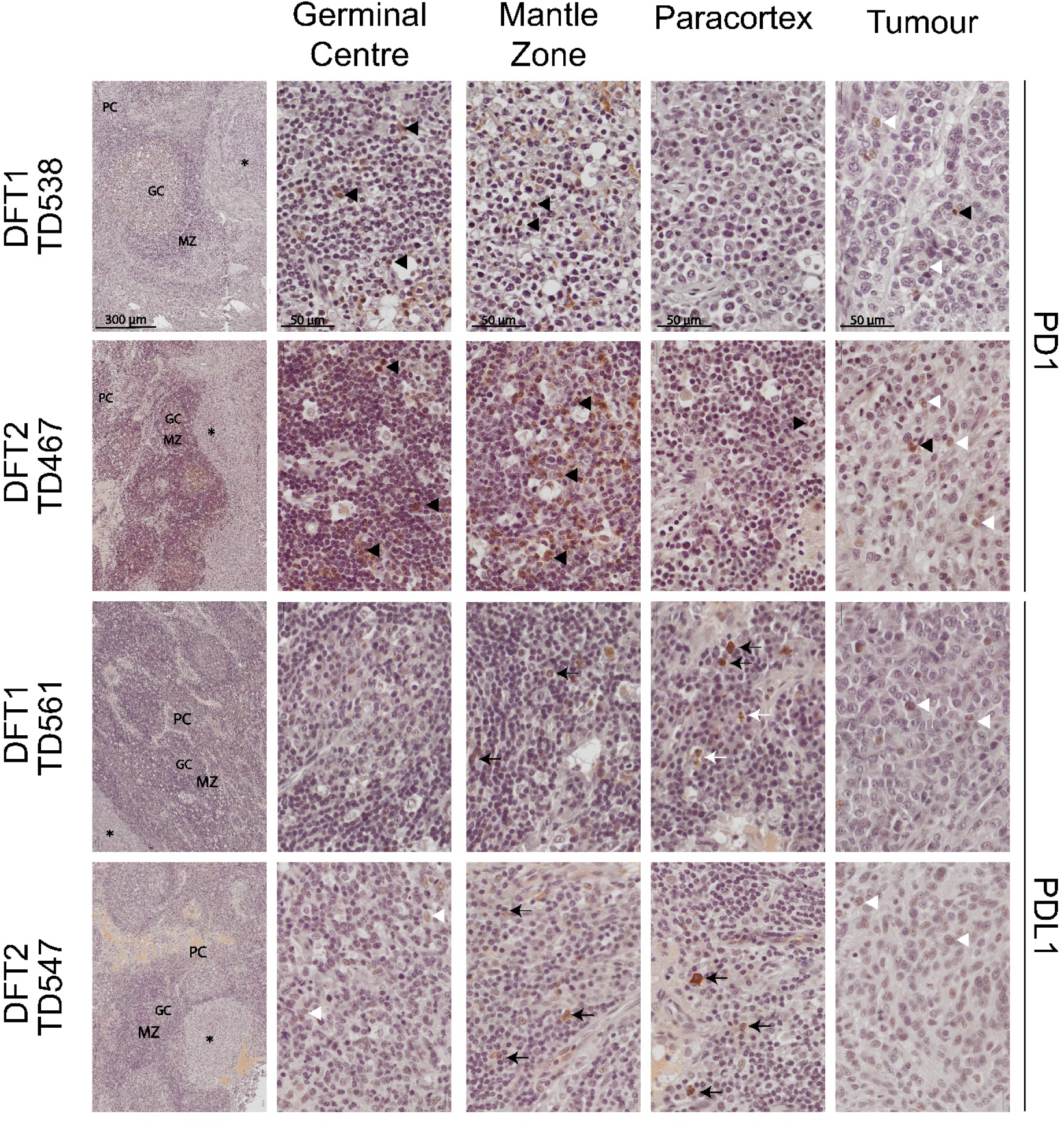
PD1 and PDL1 labeling in devil lymph nodes, DFT1 and DFT2 tumors. Representative images of DFT1 and DFT2 infected lymph nodes, and tumor metastases. PD1 membrane labeling (black arrowheads) was identified predominantly within the germinal center and mantle zone, with fewer positively labeled cells in the paracortex and tumors. PDL1 labeled cells (black arrows) were predominantly located in the paracortex. Non-specific labeling of nuclei (white arrowheads) occurred in DFT1 and DFT2 tumors, and occasional germinal centers. Hemosiderin (white arrows) was manually excluded from analysis.

Strong cytoplasmic labeling for PDL1 was observed in lymphocytes and plasma cells in all lymph node samples. PDL1 labeled cells were predominantly located in the paracortex and mantle zones (**Figure 5)**. No PDL1 labelled tumor cells were identified, however PDL1 labelled immune cells were detected at low rates (**Figure 6D**). The strength of PDL1 labeling varied between slides with DAB intensity completely obscuring the nuclei in some sections, but moderate to weak in other sections.

**Figure 6.**
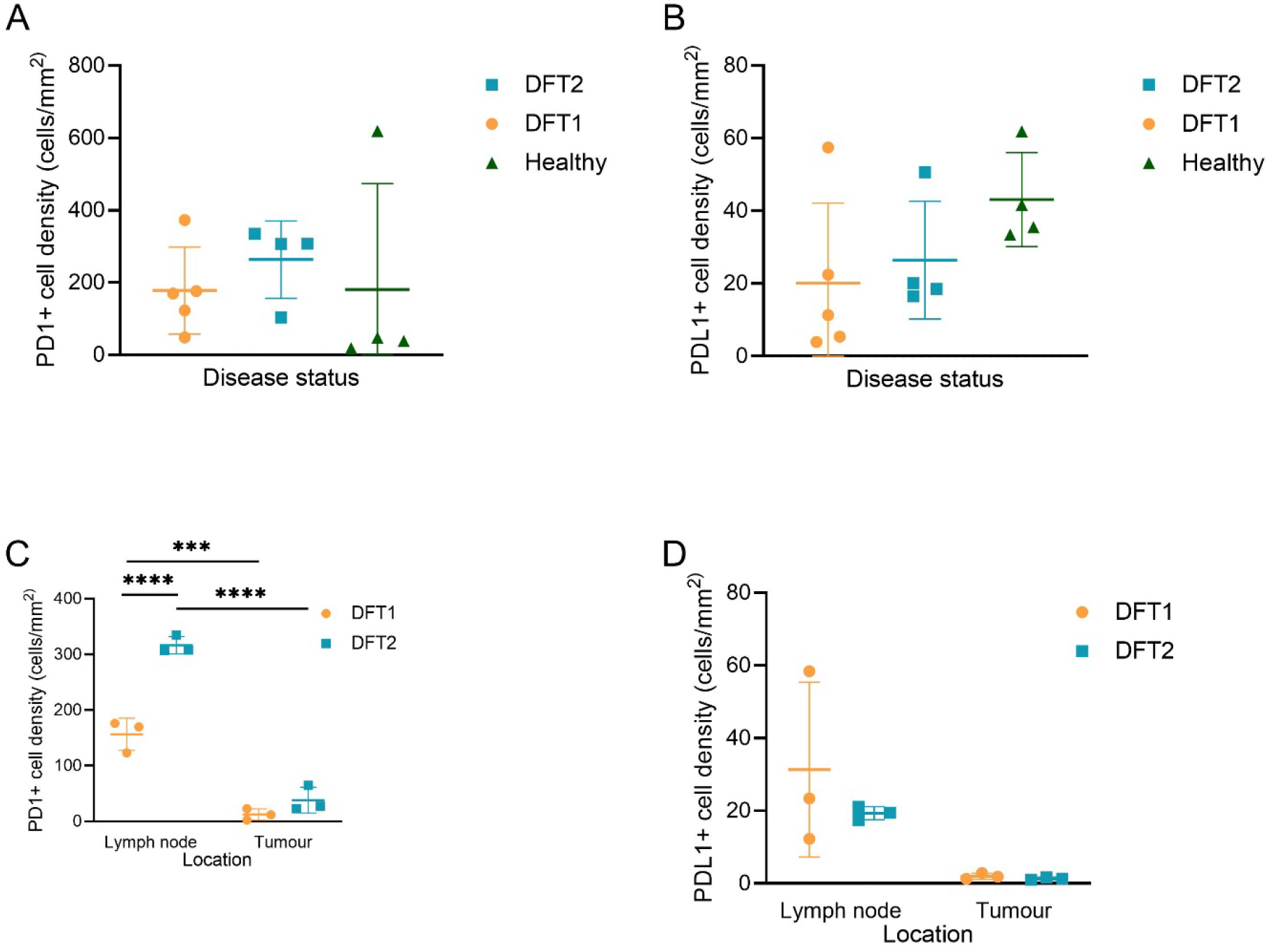
Quantification PD1 and PDL1 in lymph nodes and the tumor microenvironment. The density of PD1 and PDL1 labelled cells in devil lymph nodes was determined using the automated software QuPath. (A) PD1+ cell density in lymph nodes. Non-normally distributes data were log transformed and the difference in PD1 density assessed using a one-way ANOVA without repeated measures. No statistically significant difference was determined in PD1+ cell density between lymph nodes from healthy, DFT1 positive, or DFT2 positive devils (F(2,10) = 1.6, p = 0.25). (B) PDL1+ cell density in lymph nodes. Non-normally distributes data were log transformed and the difference in PD1 density assessed using a one-way ANOVA without repeated measures. No statistically significant difference was determined in PD1+ cell density between lymph nodes from healthy, DFT1 positive, or DFT2 positive devils (F(2,10) = 2.83, p = 0.1). (C) PD1+ cell density in lymph nodes and tumor metastases. A two-way ANOVA with Tukey multiple comparisons test revealed there was a statistically significant interaction between the presence of a metastatic DFT1 or DFT2 tumor and the PD1+ expression in lymph nodes (F(1,8) = 31.35, p = 0.0005). Lymph nodes containing DFT2 tumor metastases had a significantly higher density of PD1+ cells when compared to those with DFT1 metastases (p<0.0001). (D) PDL1+ cell density in lymph nodes and tumor metastases. A two-way ANOVA revealed there was no statistically significant interaction between the presence of a metastatic DFT1 or DFT2 tumor and the PDL1+ expression in lymph nodes (F(1,8) = 0.67, p =0.44). Error bars represent standard deviation. **p< 0.01, ****p<0.0001.

### Quantification of PD1 and PDL1 expression in lymph nodes

There was no significant difference in the density of PD1 (**Figure 6A**) or PDL1 (**Figure 6B**) expressing cells in the lymph nodes between healthy, DFT1, and DFT2, infected devils (**Table S6**). The PD1 positive cell density for TD608 was 619 cells/mm^2^, the highest density of all devils included in the study. This devil was the only wild control devil and had a higher number of secondary follicles compared to the other healthy (captive) devils (**Figure 6A**).

Lymph nodes containing tumor metastases (DFT1 n = 3; DFT2 n = 3) were sub-selected for further analysis. Lymph nodes were defined as the normal lymphoid tissue excluding tumor metastases. Tumors were defined as the whole tumor metastases internal from the fibrous capsule (DFT1) or the junction of DFT2 and normal lymph node cells (DFT2). There was a significant increase in the density of PD1 expressing immune cells within lymph nodes containing DFT2 metastatic tumors, compared to those with DFT1 metastatic tumors (Figure 6C, p <0.0001). There was no significant difference in the density of PDL1 expression between the DFT1 and DFT2 cohorts (Figure 6D).

When comparing the density of PD1 and PDL1 between lymph node and tumor metastasis, we found significantly more PD1 labelled cells in the lymph node compared to the tumor metastasis in both DFT1 and DFT2 (**Figure 6C-D**).

## Discussion

The identification and quantification of spatial relationships among cells and molecules is an essential component of immunology research. This has traditionally been assessed using light microscopy and subjective scoring. Here, a semi-automated PD1 and PDL1 scoring approach was designed using the software QuPath, and tested in comparison to manual quantification, to assess its potential use as a tool for research comparative immunology. Results from automated nuclei detection scripts, cell classifier training outcomes, and manual counting were highly correlated and highly specific, precise, and accurate. These outcomes suggest the automated image analysis strategy we developed is an appropriate alternate approach for high-throughput quantification, compared with manual counting.

Automated detection of PDL1 labelled tissue was complicated by strong cytoplasmic DAB labelling, which often obscured nuclei thereby omitting them from detection. Additionally, accurate detection of DFTD metastases was complicated by the rigid nature of nuclei detection used in this workflow. In these cases, cell detection scripts were visually assessed, and appropriate scripts were selected that accounted for both correlation and correct identification of positively labelled cells. The cell classifiers produced in this project successfully identified positively labelled cells in both PD1 and PDL1 labelled tissue (i.e., were specific). However, the classifiers tended to under-estimate the number of positively labeled cells leading to false negatives. This is not necessarily an unfavorable outcome, as consistent false negatives are more desirable than false positives. To overcome issues such as background labelling interfering with automated classifier performance, future studies would benefit from antibodies that have clear cytoplasmic and membrane staining, rather than variable patterns than can obscure the nucleus, as we observed in this work. Furthermore, as automated cell quantification technologies improve, the impact that variable cellular and nuclei staining and anatomy has on performance of automated systems will reduce (44).

When the aim of a project is absolute quantification of positive cells in a tissue section, automated image analysis offers a distinct time advantage over manual counting. Project setup and training are primarily dependent on the length of time taken to annotate tissue sections. The manual nuclei annotation steps performed in this project required approximately 30 minutes per image. The time taken to optimize nuclei detection and classifier training for each image was between two and three hours. In comparison, the *absolute quantification* of positive cells in an entire tissue section by manual annotation was estimated to take upwards of 24 hours. Thus, studies that require analysis of many slides expected to have similar tissue architecture and staining patterns will be more efficient using automated analysis. However, the outcomes achieved with QuPath, and other image analysis software are user dependent. Expert knowledge is needed to train and assess algorithms, but when used appropriately can broaden information acquired from immunohistochemically stained tissue section and increase throughput following algorithm optimization.

We observed no significant difference in the number of PD1 and PDL1 labelled cells in healthy devils compared with those with DFT1 or DFT2. This analysis was however complicated by large individual variations within each group, which we discuss in detail below. A planned sub- group analysis of the density of PD1 and PDL1 positive cells in lymph nodes containing DFT1 or DFT2 metastases revealed a significant increase in PD1 expression in the lymph nodes of devils with DFT2 metastases.

DFT1 cells are rarely observed to express MHC-I in samples from wild devils while DFT2 cells constitutively express MHC-I (11,19). MHC-I molecules present on DFT2 cells may provide antigenic stimulation which could lead to an increased reliance on alternative tumor immune evasion pathways, such as PD1/PDL1. The high expression of PD1 in germinal centers of lymph nodes containing DFT2 metastasis suggests that ample targets are available for potential anti-PD1 immune checkpoint therapy, as seen in human patients with elevated PD1 expression in lymph nodes (25).

PD1 labeling within lymph nodes is predominantly identified in germinal centers (45). Antigenic stimulation in the lymphatic drainage region directly stimulates activity within that node, resulting in an increased number of germinal centers (46,47). The variable degrees of inflammation caused by DFTD tumors, necrosis or infection of tumor tissue, and other diseases may have contributed to increases of PD1 density in some devils. Other causes of lymphoid hyperplasia associated with non-specific antigenic stimulation cannot be ruled out.

Metastases are present in most cases of DFT1 and DFT2, but little is known about the metastatic process of transmissible cancers. Ongoing transmission of tumor cells requires external tumor cells that can be transmitted through biting, so metastatic tumor cells are an evolutionary dead end for transmissible cancers. However, in humans, tumor colonization of tumor draining lymph nodes results in immune tolerance of tumor cells and facilitate distant metastasis (48,49). General immune tolerance or suppression by DFT1 or DFT2 cells could facilitate growth of multiple facial tumors and thus increase the probability of transmission to new hosts. Future studies should prioritize collection of tumor-draining lymph nodes and more distant lymph nodes to assess for inhibitory checkpoint molecules such as PDL1 and CD200 (50).

Lymph node anatomy was impacted by the presence of tumor metastases within the node. Metastatic tumor location and size within the lymph nodes was inconsistent and tumor infiltration may have altered the relative size of the medulla or cortex, thus altering the relative density of germinal centers and PD1 positive immune cells. It would be worthwhile exploring the differences observed in this study further, as they may lead to a better understanding of the differences in pathophysiology and therefore potential unique treatments for DFT1 and DFT2.

An unexpected observation was the strikingly low numbers of secondary follicles and density of PD1 expression in the lymph nodes of captive devils when compared to the wild devil without signs of DFT1 or DFT2. A difference between lymphoid tissue of captive and wild devils has not been remarked on in previous studies (51,52) and the possibility of an early stage DFT1 or DFT2 infection or exposure to other antigens cannot be ruled out. Further investigation into the immune system of captive devils would be worthwhile and is particularly important given their role as an insurance population that could be used to supplement the wild devil population and guard against extinction. While it would be premature to draw any conclusions regarding the immune landscape of captive devils through this observation, it does reflect similar findings in other wild and captive animals (53–56).

Beura et al. (54) found that adult laboratory mice housed in standard disease-free conditions did not display a mature immune phenotype. However, alterations to the environment of these mice (through co-housing with wild caught mice) resulted in maturation of the immune system. Striking differences in immune responses between wild and captive animals have also been observed in other species (55,57,58). For example, Flies et al., (55) found the serum antibody concentrations of captive non-domesticated spotted hyenas (*Crocuta crocuta)* housed in a non-sterile facility were significantly lower than those of wild hyenas. Likewise, the captive devils in this study were housed in outdoor enclosures and fed a carrion diet intended to resemble conditions of their wild counterparts. Despite this, we found a notable difference in lymph node structure between the healthy captive and wild devils.

## Conclusions

Our results suggest that PD1/PDL1 interactions are unlikely to be a primary immune evasion pathway exploited by DFT1/2 cells. However, expression patterns of PD1 and PDL1 in lymph nodes and weakly immunogenic tumors are similar and the high expression of PD1 in lymph nodes suggests the potential for broad activation of lymphocytes via PD1 blockade. This information may assist with the development of prophylactic and therapeutic strategies for DFT1 and DFT2 (59,60). The semi-automated whole image analysis pipeline developed for this project produced high quality results and could play an important role in analyzing anti-tumor responses in large scale field trials. Additionally, understanding the role of immune checkpoints in transmissible cancers and allograft immune evasion in devils may provide additional information for other cancers and inform fields such as transplant medicine.

## Abbreviations

PD1: programmed cell death protein 1
PDL1: programmed death ligand 1
DFT1: devil facial tumor 1
DFT2: devil facial tumor 2
DFTD: devil facial tumor disease
MCC: Matthew’s correlation coefficient;
DAB: diaminobenzidine
ROI: region of interest

## Acknowledgements

We thank Dr Graeme Knowles from the Department of Natural Resources and Environment, Tasmania, for guidance on the histological assessment of DFTD and the generous provision of healthy Tasmanian devil tissue used in this study. We thank Dr Jo-Maree Courtney for their valuable assistance in writing scripts for use in QuPath.

## Author contributions

GGR: conceptualization, methodology, validation, formal analysis, investigation, writing – original draft, writing – review and editing, visualization. AF: conceptualization, methodology, validation, formal analysis, investigation, writing – original draft, writing – review and editing, visualization, supervision, funding acquisition, resources. GM: methodology, writing – review and editing. RP: writing – review and editing, supervision. CP: resources, supervision, writing – review and editing. NFJ: formal analysis, methodology. JD: methodology, writing – review and editing.

## Data availability

Data including raw images available upon request.

