## Supplementary Materials for "Automated analysis of PD1 and PDL1 in lymph nodes and the microenvironment of transmissible tumors in Tasmanian devils"

Andrew S. Flies, PhD

Menzies Institute for Medical Research, College of Health and Medicine

University of Tasmania

Private Bag 23, Hobart TAS 7000

#### Supplementary Methods 1

#### Processing of Tasmanian devil samples

### 1 **Tasmanian devil tissue origin and processing**

DFT1 and DFT2 tumor sections and devil lymphoid tissue samples were sourced from paraffin embedded samples supplied by the tissue bank at Menzies Institute for Medical Research, University of Tasmania. One healthy (non-DFTD) devil lymph node was sourced from a wild devil found as roadkill. It was found within 2-3 hours of death and stored, on ice, for approximately 12 hours prior to necropsy. The three remaining healthy lymph node samples were provided by the Department of Natural Resources and Environment Tasmania (NRET) Tasmania sourced from healthy captive devils. Information on the sex of these healthy captive devil was not provided.

### 2 **Processing and paraffin embedding of tissue samples**

Samples from TD467, TD500	, TD504, TD510, TD538, TD547, TD549, TD561, TD588, and TD608 had been formaldehyde fixed and paraffin embedded as part of previous studies. Tasmanian devil tissue sections from TD1296, 1297, 1298 and 1299 were collected at necropsy following euthanasia and stored in formaldehyde for at least 24 hours, before transferred being to 70% ethanol They were stored in 70% ethanol at room temperature until paraffin embedding. No record for the duration of formaldehyde fixation of ethanol storage was available. Paraffin embedding of lymph node sections was performed using a tissue auto processor (Lecia ASP 300S) on the routine overnight program, omitting the formaldehyde step, according to the manufacturer’s instructions. Paraffin embedded tissue samples were stored at room temperature until cutting.

### 3 **Cutting and mounting paraffin embedded tissue**

Paraffin blocks were trimmed at 20 μm thickness until a representative section of tissue was obtained. The paraffin block face was then polished through gentle cutting of 3-4 sections at 4 μm which were discarded. Following polishing, 4 μm sections were cut and floated on 48^o^C water bath before mounting into FLEX IHC microscope slides (K8020). Mounted sections were incubated at 37^o^C for a minimum of 12 hours to allow to dry, then stored at room temperature for a maximum of 4 weeks prior to staining.

**Table S1 Details of Tasmanian devils from which samples for analysis were obtained**

| ID | Age | Sex | Health information | Tissue sections |
| --- | --- | --- | --- | --- |
| TD1296 | <1 | NR | DFT negative. No illness | Lymph node |
| TD1297 | <1 | NR | DFT negative. No illness | Lymph node |
| TD1299 | <1 | NR | DFT negative. No illness | Lymph node |
| TD608 | 1 | M | DFT negative. Dead roadkill | Submandibular lymph node |
| TD538 | 5 | M | DFT1 | Submandibular lymph node 1 with metastases, |
| TD504 | 1 | F | DFT1 | Submandibular lymph node |
| TD510 | 1 | F | DFT1 | Lymph node |
| TD588 | 2 | M | DFT1 | Submandibular lymph node with metastases |
| TD561 | NR | M | DFT1 | Lymph node with metastases |
| TD500 | 3 | M | DFT2 | Lymph node with metastases |
| TD549 | 3 | M | DFT2 | Submandibular lymph node |
| TD547 | 3 | M | DFT2 | Submandibular lymph node |
| TD467 | 3 | M | DFT2 | Submandibular lymph node with metastases |

*DFT1* – devil facial tumor 1, *DFT2 –* devil facial tumor 2, *NR –* not recorded.

**Table S2.** Disease status and lymph node characteristics of all devils included in this investigation.

| Lymph node characteristics | | | | | |
| --- | --- | --- | --- | --- | --- |
| Devil ID | Disease status | Lymph node area – ex. tumor metastases (mm^2^) | Tumor metastases area (mm^2^) | No. secondary follicles | follicles/mm^2^ |
| TD1296 | Negative | 32.60 | n/a | 9.00 | 0.28 |
| TD1297 | Negative | 16.40 | n/a | 2.00 | 0.12 |
| TD1299 | Negative | 12.90 | n/a | 6.00 | 0.47 |
| TD608 | Negative (Roadkill) | 30.50 | n/a | 39.00 | 1.28 |
| TD538 | DFT1 | 135.00 | 9.90 | 140.00 | 1.04 |
| TD504 | DFT1 | 124.00 | n/a | 147.00 | 1.19 |
| TD510 | DFT1 | 122.20 | n/a | 114.00 | 0.93 |
| TD588 | DFT1 | 28.40 | 2.80 | 29.00 | 1.02 |
| TD561 | DFT1 | 27.80 | 140.90 | 20.00 | 0.72 |
| TD500 | DFT2 | 8.80 | 51.00 | 23.00 | 2.61 |
| TD549 | DFT2 | 72.00 |  | 25.00 | 0.35 |
| TD547 | DFT2 | 122.10 | 0.30 | 216.00 | 1.77 |
| TD467 | DFT2 | 48.80 | 9.90 | 50.00 | 1.02 |

**Table S3.** Densities of PD1/PDL1 in DFT1/DFT2/healthy devil lymph nodes and tumor metastases (if present) in lymph nodes with and without tumor metastases

|  | DFT1 | | | | | DFT2 | | | | Healthy | | | |
| --- | --- | --- | --- | --- | --- | --- | --- | --- | --- | --- | --- | --- | --- |
| Devil ID | TD538 | TD588 | TD561 | TD504 | TD510 | TD547 | TD500 | TD467 | TD549 | TD608 | TD1296 | TD1297 | TD1299 |
| PD1 positive cell density (cells/mm2) | | | | | | | | | | | | | |
| Lymph node | 176.12 | 169.90 | 123.16 | 372.87 | 48.56 | 307.73 | 307.11 | 334.74 | 103.77 | 619.16 | 47.98 | 18.23 | 38.57 |
| Tumor | 2.66 | 22.79 | 11.75 | n/a | n/a | 64.89 | 27.09 | 22.68 | n/a | n/a | n/a | n/a | n/a |
| PDL1 positive cell density (cells/mm2) | | | | | | | | | | | | | |
| Lymph node | 11.29 | 57.41 | 22.42 | 3.88 | 5.39 | 18.47 | 20.08 | 16.48 | 50.60 | 33.45 | 61.83 | 41.62 | 35.51 |
| Tumor | 1.91 | 0.88 | 0.28 | n/a | n/a | 0 | 0.72 | 0.39 | n/a | n/a | n/a | n/a | n/a |
| Analysis performed on each cohort (PD1 and PDL1) | | | | | | | | | | | | | |
| Lymph node: DFT1 v DFT2 v healthy | | | | | | | | | | | | | |
| Tumor: DFT1 v DFT2 | | | | | | | | | | | | | |
| Lymph node (with metastases): DFT2 V DFT1 | | | | | | | | | | | | | |
| DFT1: LN V Tumor | | | | | | | | | | | | | |
| DFT 2: LN V tumor | | | | | | | | | | | | | |
| Definitions of regions | | | | | | | | | | | | | |
| Lymph node: lymph node excluding whole tumor metastases | | | | | | | | | | | | | |
| Tumor: whole tumor metastases, internal from the fibrous capsule (DFT1) or the junction of DFT2 and lymph cells (DFT2) | | | | | | | | | | | | | |

**Table S4.** Densities of PD1/PDL1 in DFT1 and DFT2 lymph nodes *with* tumor metastases divided into: Lymph node, tumor outer margin, tumor inner margin, and tumor center.

|  | DFT1 | | |  | DFT2 | | |
| --- | --- | --- | --- | --- | --- | --- | --- |
| Devil ID | TD538 | TD588 | TD561 |  | TD547 | TD500 | TD467 |
| PD1 positive cell density (cells/mm^2^) | | | | | | | |
| Lymph node | 176.1 | 169.6 | 155.7 |  | 301.8 | 473.0 | 301.7 |
| Tumour outer margin | 140.2 | 171.1 | 74.6 |  | 735.4 | 215.9 | 625.4 |
| Tumour Inner Margin | 4.4 | 25.1 | 22.3 |  | 64.8 | 36.6 | 25.8 |
| Tumour centre | 0.4 | 8.0 | 9.9 |  | 0 | 23.0 | 0 |
| PDL1 positive cell density (cells/mm^2) | | | | | | | |
| Lymph node | 11.0 | 54.9 | 25.4 |  | 18.4 | 29.6 | 17.3 |
| Tumour outer margin | 18.0 | 69.1 | 17.3 |  | 20.7 | 13.7 | 9.0 |
| Tumour Inner Margin | 3.5 | 0.9 | 0.9 |  | 0 | 1.5 | 0.4 |
| Tumour centre | 0.1 | 0 | 0.1 |  | 0 | 0.3 | 0.2 |
| **Analysis performed on each cohort (PD1 and PDL1):** | | | | | | | |
| Tumour centre: DFT1 v DFT2 | |  |  |  |  |  |  |
| Tumour Inner Margin DFT1 v DFT2 | | |  |  |  |  |  |
| Tumour outer margin: DFT1 v DFT2 v healthy lymph node | | | | | |  |  |
| DFT1: tumour inner margin v tumour centre v lymph node V tumour outer margin | | | | | | | |
| DFT 2: tumour inner margin v tumour centre v lymph node v tumour outer margin | | | | | | | |
| **Definitions of regions** | |  |  |  |  |  |  |
| Lymph node: anatomically normal lymph node excluding tumour metastases and tumour outer margin | | | | | | | |
| Tumour outer margin: 500um margin from the edge of the tumour into the lymph node | | | | | | | |
| Tumour inner margin: 500um margin from the edge of the tumour metastases internally | | | | | | | |
| Tumour centre: centre of the tumour from the 500um margin. | | | | | |  |  |

|  | | **Classification of cell based off ‘ground truth’ (manual counting)** | | |
| --- | --- | --- | --- | --- |
|  |  | Positive | Negative | Measurements |
| **Result from automated test** | Positive | True Positive (TP) | False Positive (FP) | *Precision* $\frac{\mathrm{TP}}{TP+FP}$ |
|  | Negative | False Negative (FN) | True Negative (TN) | *Negative predictive value* $\frac{\mathrm{TN}}{TN+FN}$ |
|  | Measurements | *Sensitivity* $\frac{\mathrm{TP}}{TP+FN}$ | *Specificity* $\frac{\mathrm{TN}}{TN+FP}$ | *Accuracy*$\frac{TP+TN}{TP+TN+FP+FN}$ |

**Figure S1 Confusion matrix used to assess cell classification methods.** A confusion matrix was constructed using the true positive (TP), true negative (TN), false positive (FP), and false negative (FN) measurements for each cell classification method. The outcomes of the confusion matrix could then be used to compare cell classification methods.

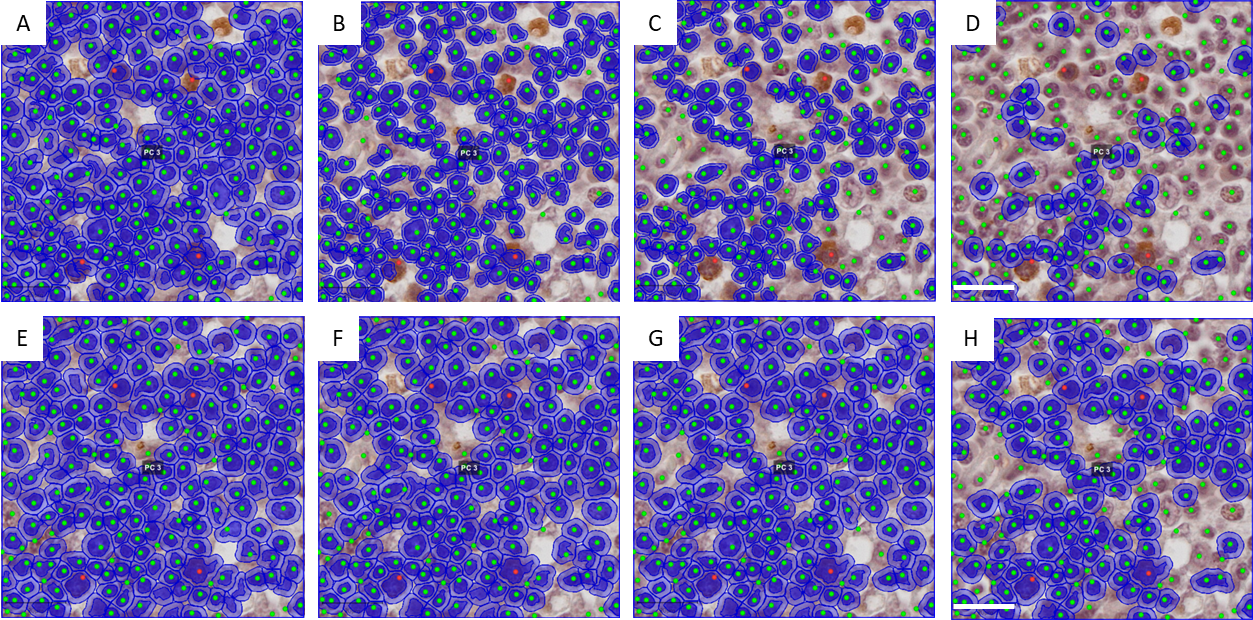

**Figure S2.** Nuclei detection performed using various cell detection algorithms. Nuclei detection using haematoxylin optical density was used in scripts 1, 4, 8, 12 (A-D respectively) demonstrating identification of nuclei cells while excluding some cells with strong cytoplasmic DAB staining. Optical density sum was used for nuclei detection in scripts 13, 17, 20, 24 (E-H respectively), demonstrating identification of cells with strong cytoplasmic DAB staining. The threshold of cell detection parameters was increased between A-D and E-F which is reflected with the decrease in the number of nuclei identified. The correlations between the manual nuclei counts and all automated nuclei detection scripts were then assessed to select the most appropriate settings for the image. TD500 lymph node, anti-PDL1 (1F8), 40x magnification. Scale bar 20 μm.

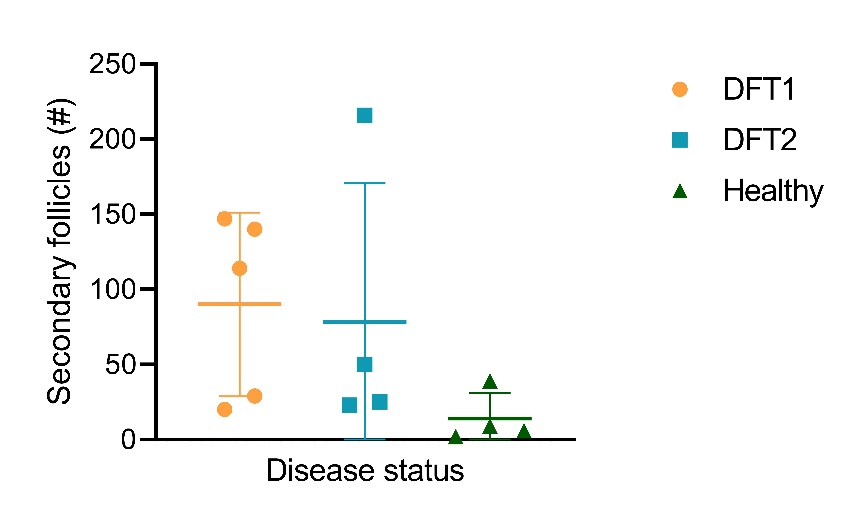

**Figure S3.** Secondary follicles in healthy, DFT1 and DFT2 infected Tasmanian devil lymph nodes. A Kruskal-Wallis test revealed there was no statistically significant different in the number of secondary follicles in lymph nodes from healthy, or DFT1 or DFT2 infected Tasmanian devils (Kruskal-Wallis statistic = 4.711, p=0.0924).

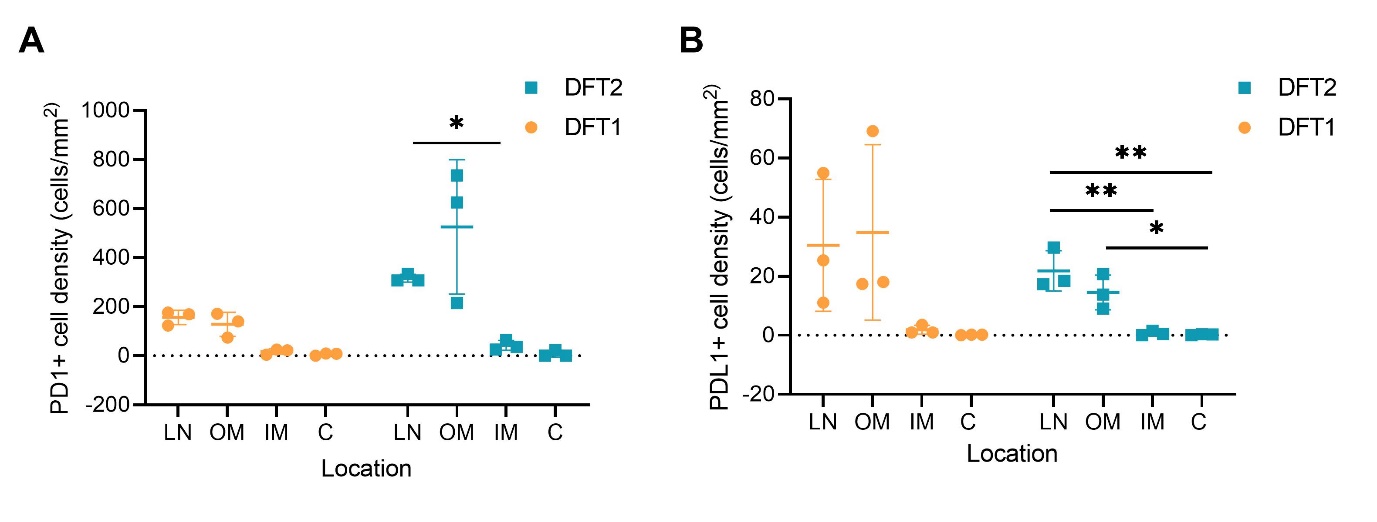

**Figure S4.** **Quantification PD1 and PDL1 in lymph nodes containing metastatic tumors.** The density of PD1 and PDL1 positive cells in devil lymph nodes was determined using the automated software QuPath. PD1+ (A) and PDL1+ (B) in the metastatic tumor microenvironment. Two-way ANOVAs with repeated measures and Tukey’s multiple comparison were used to assess the density of PD1+ (C) and PDL1+ (D) cells in lymph nodes and the outer margin (OM), inner margin (IM), and center (CE) of the tumor microenvironment. Šídák's multiple comparisons test were used to compare DFT1 and DFT2 in (C) and (D). Error bars represent standard deviation. *p<0.05, **p< 0.01. LN – lymph node, OM- tumor outer margin, IM – tumor inner margin, C – tumor center**.**
