## Supplementary dataset 2 for "Automated analysis of PD1 and PDL1 in lymph nodes and the microenvironment of transmissible tumors in Tasmanian devils": Cell Detection V13.groovy

```
clearDetections()
selectAnnotations()
runPlugin('qupath.imagej.detect.cells.WatershedCellDetection',
        '{"detectionImageBrightfield": "Optical density sum",  ' +
                '"requestedPixelSizeMicrons": 0.1,  ' +
                '"backgroundRadiusMicrons": 0.0,  "medianRadiusMicrons": 0.0,  ' +
                '"sigmaMicrons": 1.0,  ' +
                '"minAreaMicrons": 10.0,  ' +
                '"maxAreaMicrons": 400.0,  ' +
                '"threshold": 0.1,  ' +
                '"maxBackground": 2.0,  ' +
                '"watershedPostProcess": true,  ' +
                '"excludeDAB": false,  ' +
                '"cellExpansionMicrons": 2.0,  ' +
                '"includeNuclei": true,  ' +
                '"smoothBoundaries": true,  ' +
                '"makeMeasurements": true}');
print "done!"
```
