## Supplementary dataset 2 for "Automated analysis of PD1 and PDL1 in lymph nodes and the microenvironment of transmissible tumors in Tasmanian devils": Supplementary dataset 2.docx

**Supplementary dataset 2: nuclei detection scripts**

Cell detection scripts were produced in intelliJ community edition.

27 scripts were produced in total, however only 24 of these were used for the study as there were errors in script numbers 18, 19, and 23.

These scripts and their results were removed from the study.

Scripts were renamed as below:

| Original name | New name |
| --- | --- |
| Cell Detection V1 | Cell Detection V1 |
| Cell Detection V2 | Cell Detection V2 |
| Cell Detection V3 | Cell Detection V3 |
| Cell Detection V4 | Cell Detection V4 |
| Cell Detection V5 | Cell Detection V5 |
| Cell Detection V6 | Cell Detection V6 |
| Cell Detection V7 | Cell Detection V7 |
| Cell Detection V8 | Cell Detection V8 |
| Cell Detection V9 | Cell Detection V9 |
| Cell Detection V10 | Cell Detection V10 |
| Cell Detection V11 | Cell Detection V11 |
| Cell Detection V12 | Cell Detection V12 |
| Cell Detection V13 | Cell Detection V13 |
| Cell Detection V14 | Cell Detection V14 |
| Cell Detection V15 | Cell Detection V15 |
| Cell Detection V16 | Cell Detection V16 |
| Cell Detection V17 | Cell Detection V17 |
| Cell Detection V20 | Cell Detection V18 |
| Cell Detection V21 | Cell Detection V19 |
| Cell Detection V22 | Cell Detection V20 |
| Cell Detection V24 | Cell Detection V21 |
| Cell Detection V25 | Cell Detection V22 |
| Cell Detection V26 | Cell Detection V23 |
| Cell Detection V27 | Cell Detection V24 |
