## Supplementary dataset 2 for "Automated analysis of PD1 and PDL1 in lymph nodes and the microenvironment of transmissible tumors in Tasmanian devils": tumour_margin.groovy

```
import org.locationtech.jts.geom.Geometry
import qupath.lib.common.GeneralTools
import qupath.lib.objects.PathObject
import qupath.lib.objects.PathObjects
import qupath.lib.roi.GeometryTools
import qupath.lib.roi.ROIs

import static qupath.lib.gui.scripting.QPEx.*


// How much to expand each region
double expandMarginMicrons = 500.0

// Define the colors
def coloInnerMargin = getColorRGB(0, 0, 200)
def colorOuterMargin = getColorRGB(0, 200, 0)
def colorCentral = getColorRGB(0, 0, 0)

// Choose whether to lock the annotations or not (
def lockAnnotations = true

def imageData = getCurrentImageData()
def hierarchy = imageData.getHierarchy()
def server = imageData.getServer()

def cal = server.getPixelCalibration()
if (!cal.hasPixelSizeMicrons()) {
    print 'We need the pixel size information here!'
    return
}
if (!GeneralTools.almostTheSame(cal.getPixelWidthMicrons(), cal.getPixelHeightMicrons(), 0.0001)) {
    print 'Warning! The pixel width & height are different; the average of both will be used'
}

// Get annotation & detections
def annotations = getAnnotationObjects()
def selected = getSelectedObject()
if (selected == null || !selected.isAnnotation()) {
    print 'Please select an annotation object!'
    return
}
annotations.remove(selected)

// Extract the ROI & plane
def roiOriginal = selected.getROI()
def plane = roiOriginal.getImagePlane()

// If we have at most one other annotation, it represents the tissue
Geometry areaTissue
PathObject tissueAnnotation
if (annotations.isEmpty()) {
    areaTissue = ROIs.createRectangleROI(0, 0, server.getWidth(), server.getHeight(), plane).getGeometry()
} else if (annotations.size() == 1) {
    tissueAnnotation = annotations.get(0)
    areaTissue = tissueAnnotation.getROI().getGeometry()
} else {
    print 'Sorry, this script only support one selected annotation for the tumor region, and at most one other annotation to constrain the expansion'
    return
}

// Calculate how much to expand
double expandPixels = expandMarginMicrons / cal.getAveragedPixelSizeMicrons()
def areaTumor = roiOriginal.getGeometry()

// Get the outer margin area
def geomOuter = areaTumor.buffer(expandPixels)
geomOuter = geomOuter.difference(areaTumor)
geomOuter = geomOuter.intersection(areaTissue)
def roiOuter = GeometryTools.geometryToROI(geomOuter, plane)
def annotationOuter = PathObjects.createAnnotationObject(roiOuter)
annotationOuter.setName("Outer margin")
annotationOuter.setColorRGB(colorOuterMargin)

// Get the central area
def geomCentral = areaTumor.buffer(-expandPixels)
geomCentral = geomCentral.intersection(areaTissue)
def roiCentral = GeometryTools.geometryToROI(geomCentral, plane)
def annotationCentral = PathObjects.createAnnotationObject(roiCentral)
annotationCentral.setName("Center")
annotationCentral.setColorRGB(colorCentral)

// Get the inner margin area
def geomInner = areaTumor
geomInner = geomInner.difference(geomCentral)
geomInner = geomInner.intersection(areaTissue)
def roiInner = GeometryTools.geometryToROI(geomInner, plane)
def annotationInner = PathObjects.createAnnotationObject(roiInner)
annotationInner.setName("Inner margin")
annotationInner.setColorRGB(coloInnerMargin)

// Add the annotations
hierarchy.getSelectionModel().clearSelection()
hierarchy.removeObject(selected, true)
def annotationsToAdd = [annotationOuter, annotationInner, annotationCentral];
annotationsToAdd.each {it.setLocked(lockAnnotations)}
if (tissueAnnotation == null) {
    hierarchy.addPathObjects(annotationsToAdd)
} else {
    tissueAnnotation.addPathObjects(annotationsToAdd)
    hierarchy.fireHierarchyChangedEvent(this, tissueAnnotation)
    if (lockAnnotations)
        tissueAnnotation.setLocked(true)
}
```
